# Trophic strategies explain the ocean niches of small eukaryotic phytoplankton

**DOI:** 10.1101/2022.02.27.482152

**Authors:** Kyle F. Edwards, Qian Li, Kelsey A. McBeain, Christopher R. Schvarcz, Grieg F. Steward

**Affiliations:** Department of Oceanography, School of Ocean and Earth Science and Technology (SOEST), University of Hawai‘i at Mānoa; Honolulu, Hawai‘i, 96822, USA; Daniel K. Inouye Center for Microbial Oceanography: Research and Education, School of Ocean and Earth Science and Technology (SOEST), University of Hawai‘i at Mānoa; Honolulu, Hawai‘i, 96822, USA; School of Oceanography, Shanghai Jiao Tong University; China, Shanghai Shi, Xuhui Qu, 1954 Huashan Rd, 200240

## Abstract

A large fraction of marine primary production is performed by diverse small protists, and many of these phytoplankton are mixotrophs that also consume bacterial prey. However, the mechanisms structuring this diversity and its biogeochemical consequences remain poorly understood. Here we use isolates from seven major taxa to demonstrate tradeoffs between phototrophic and phagotrophic abilities. We then show that trophic strategy along the autotrophy-mixotrophy spectrum correlates strongly with global niche differences, across depths and across gradients of stratification and chlorophyll *a*. A model of competition shows that community shifts can be explained by greater fitness of faster-grazing mixotrophs when nutrients are scarce and light is plentiful. Our results illustrate how basic physiological constraints and principles of resource competition can organize complexity in the surface ocean ecosystem.

## Introduction

Photosynthesis is the foundation of Earth’s ecosystems and half of the daily primary production on the planet occurs in the surface ocean (*1*). Most of this marine primary production is carried out by single-celled phytoplankton from a broad spectrum of ancient evolutionary lineages. Many eukaryotic members of the phytoplankton live a dual lifestyle as phagotrophic mixotrophs, meaning they photosynthesize but also consume prey (*2*). The functional capabilities of many phytoplankton taxa are not known in detail, and the conditions that select for autotrophic vs. mixotrophic strategies are not well established. This is particularly true for eukaryotic phytoplankton smaller than ~5 *μ*m, which rival cyanobacteria as key photosynthesizers in the extensive oligotrophic ocean (*3–5*), while also being major predators of bacteria (*6–8*). These organisms come from many deeply diverging lineages, and encompass a rich diversity that exceeds that of smaller picocyanobacteria or larger microphytoplankton. Molecular surveys show co-occurrence of many higher taxa, such as haptophytes, chlorophytes, chrysophytes, dictyochophytes, pelagophytes, cryptophytes, bolidophytes, chlorarachniophytes, dinoflagellates, and diatoms (*9–11*). Habitat differences among clades suggest functional diversity (*12–14*), but understanding how key traits vary is hampered by a paucity of experiments on isolates, especially those cultivated from the open ocean (*3, 15, 16*). Trophic strategy may be an important axis of divergence: most taxa include phagotrophic mixotrophs that make their own chloroplasts (*6, 17*), and feeding rates vary across clades (*17*), but groups such as non-flagellated prasinophytes likely cannot ingest prey, and this may be true of some flagellates as well (*18, 19*). Thus there is likely a spectrum of trophic strategies, ranging from strictly autotrophic to largely heterotrophic phytoplankton, but data on how key functions vary across species and communities is lacking. Models predict that mixotrophy in an ecosystem should generally increase primary production, trophic transfer efficiency, and carbon export, while decreasing nutrient remineralization (*20, 21*). Therefore the prevalence of mixotrophy across ocean habitats, and the traits of the dominant taxa, may have broad consequences.

The relative fitness conferred by a given trophic strategy will depend on resource competition and the tradeoffs that constrain trait evolution. Photosynthesizers that consume prey benefit from multiple sources of nutrients and energy, but also experience competition from multiple directions, competing with specialized autotrophs for dissolved nutrients and light, and with specialized heterotrophs for prey (*22*). Mixotrophs must allocate biomass and energy among a greater number of functions than specialists, which should reduce mass-specific photosynthesis compared to autotrophs. Likewise, they may ingest prey more slowly than heterotrophs if they invest less in phagotrophy, and the operation of multiple trophic modes could increase respiratory demand (*23, 24*). However, quantification of such tradeoffs is limited, and trait comparisons have focused mostly on larger coastal dinoflagellates (*25*) and some chrysophytes (*23, 26*). For example, mixotrophic dinoflagellates tend to have lower maximal ingestion rates than similar-sized heterotrophic dinoflagellates (*25*), and they also grow more slowly than similar-sized autotrophic diatoms if dissolved nutrients and light are the only resources (*27*). It is unknown whether similar tradeoffs constrain the broader array of mixotrophic phytoplankton in open-ocean ecosystems.

Tradeoffs among trophic strategies should cause community structure to vary in predictable ways across environmental gradients. Compared to similar-sized autotrophs, phagotrophic mixotrophs should have a competitive advantage when dissolved nutrients are scarce relative to nutrients available in prey, or when light energy is limited relative to the chemical energy that can be derived from prey (*22*). At the same time, mixotrophs should do worse than strict heterotrophic predators under low light, because photosynthesis by mixotrophs is too low to compensate for their lower ingestion rates. Under high light, however, mixotrophs are expected to outperform the heterotrophs, because the energy subsidy from photosynthesis should allow them to suppress prey to densities too low to sustain heterotrophs (*28*). Therefore, the fitness of a mixotrophic strategy depends on relative supply of different resources as well as key tradeoffs, which combine to determine the net outcome of competition with multiple specialists.

Models using reasonable assumptions have found that well-lit environments with low nutrient supply may be most favorable for mixotrophs (*29, 30*), but critical physiological parameters remain poorly constrained. In a synthesis of *in situ* experiments, lower latitude environments with greater irradiance showed increasing abundance of mixotrophs relative to specialists, and mixotrophs also increased relative to heterotrophs (but not autotrophs) in nutrient-rich coastal environments, patterns which were mostly consistent with model predictions (*29*). This prior analysis considered mixotrophs and autotrophs as aggregates, but the extensive diversity within these groups raises the question of whether niche differences across taxa can be explained by trophic strategies, and whether quantifying mixotrophs in aggregate obscures important functional variation. If mixotrophs vary in their allocation of resources to different functions then the most successful strategy may vary continuously across gradients of light, nutrients, and prey (*24*). Selection for different strategies across gradients, combined with physical mixing of plankton communities, may help explain the high local diversity of small phytoplankton (*31*), while also leading to gradients in ecosystem function.

## Results

### Functional diversity and tradeoffs

In this study we first assess functional diversity and tradeoffs in a diverse guild of small open-ocean phytoplankton, and we then ask whether community variation across depth and chlorophyll *a* (Chl *a*) gradients can be explained by competition among trophic strategies. To characterize how phototrophic and phagotrophic abilities covary across species we used eleven strains of <5 *μ*m diameter eukaryotes isolated from the North Pacific Subtropical Gyre, representing eight classes that are widespread in the open ocean (Table S1, Fig. S1). Capacity for phagotrophic mixotrophy was assayed as the ability to grow with *Prochlorococcus* prey as the only added and significant source of nitrogen, under illuminated conditions. Four strains did not exhibit phagotrophic growth - two non-flagellated prasinophytes (*Ostreococcus, Chloropicon*), one flagellated prasinophyte (*Micromonas*), and one flagellated pelagophyte (*Pelagomonas*) (Fig. S2). For simplicity we will refer to these four strains functionally as ‘autotrophs’, while acknowledging the possibility that they ingest prey at very low rates that do not support growth, or could grow phagotrophically (or osmotrophically) on another food source. The remaining seven strains can grow phagotrophically (Fig. S2, (*17*)), and the rates at which they ingest *Prochlorococcus* were previously reported (*17*). To characterize how phagotrophic capacity is related to phototrophy we measured growth of all strains under phototrophic conditions (addition of dissolved nutrients but not prey) at a ‘high’ irradiance (100 *μ*mol photons m^-2^ s^-1^) that is the typical optimal irradiance for phytoplankton growth (*27*) and a ‘low’ irradiance (10 *μ*mol photons m^-2^ s^-1^) that is ~1% of surface PAR at the location from which these strains were isolated (*32*). Under low irradiance the autotrophic strains grew at rates of 0.25-0.42 d^-1^, while six of the seven mixotrophic strains failed to grow under these conditions (Fig. 1a). Under high irradiance all strains could grow phototrophically except the chrysophyte (Fig. 1a). The high irradiance growth rates of autotrophs and mixotrophs overlap, although two strains of *Florenciella* were the only mixotrophs to grow faster than the slowest-growing autotrophs. *Florenciella* exhibits relatively low ingestion rates, and across the mixotrophs the faster grazers tend to grow more slowly under high irradiance (Fig. 1b; Spearman *ρ* = −0.85, p = 0.024).

**Figure 1.**
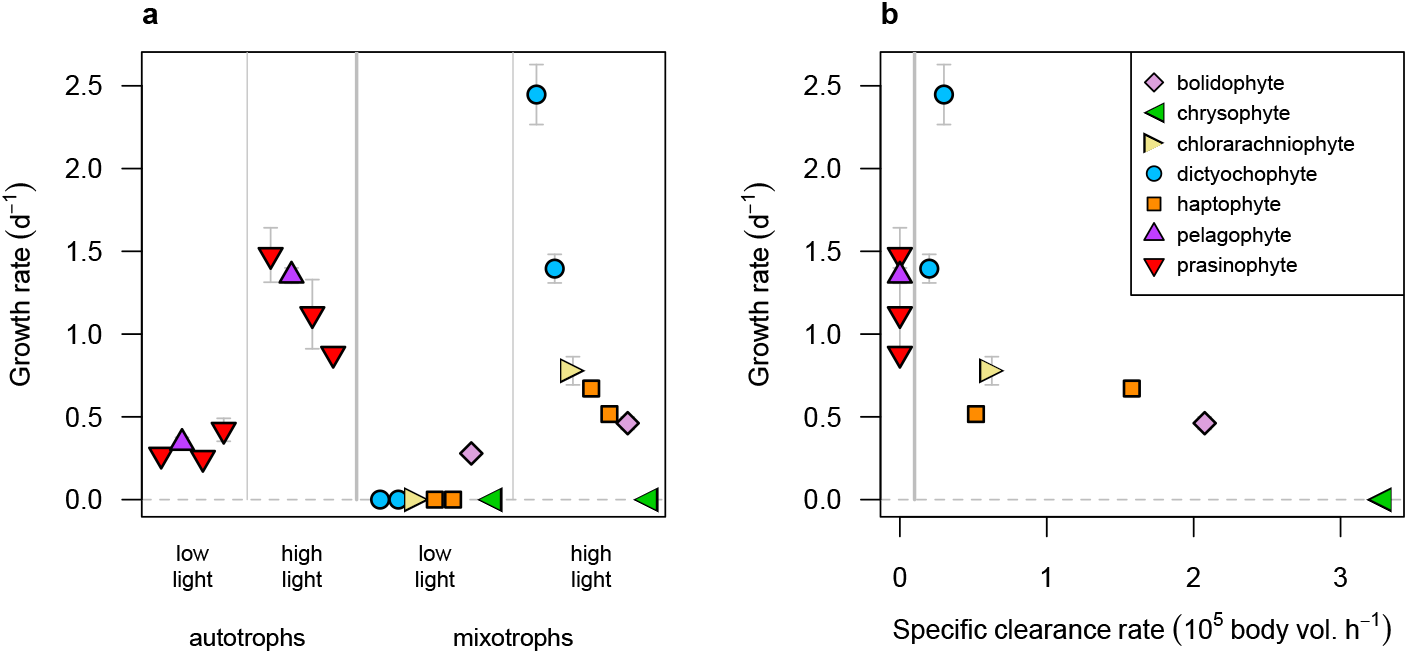
(**A**) Growth of 11 phytoplankton strains under phototrophic conditions (K medium, no added prey) when exposed to ‘high light’ (100 *μ*mol photons m^-2^ s^-1^) or ‘low light’ (10 *μ*mol photons m^-2^ s^-1^). Strains are divided into autotrophs and mixotrophs, and those exhibiting no growth are given zeros. Error bars are +/− 1 standard error of the mean; bars not visible are smaller than the point. (**B**) Growth rate under high light vs. specific clearance rate when fed *Prochlorococcus*. Strains exhibiting no ability to grow when fed prey are given zeros.

In sum these results imply that six of the mixotroph strains maintain the ability to grow photoautotrophically, but that greater grazing capacity is associated with a decline in phototrophic performance, which could be caused by reduced investment in photosynthetic machinery and/or greater respiratory demand. The cost of mixotrophy appears to be particularly high for phototrophic performance under light limitation, because only one mixotroph could grow in this treatment. The chrysophyte, which has the fastest specific maximum clearance rate of any cultivated flagellate (*17*), may be an obligate mixotroph, as it did not grow when illuminated without added prey.

### Relationship between trophic strategies and ocean niches

We next asked whether the trophic strategies of phytoplankton can explain their niches in the ocean, by combining (1) our assays of autotroph/mixotroph status; (2) our previously reported measurements of *Prochlorococcus* clearance rates for 29 mixotroph isolates (Table S2); and (3) an analysis of environmental distributions using *Tara* Oceans metabarcoding data for pico/nanoeukaryotes (size fraction 0.8-5 *μ*m) at 39 ocean stations. The *Tara* Oceans stations are primarily open ocean sites, with oligotrophic or mesotrophic characteristics (median surface Chl *a*: 0.16 *μ*g L^-1^; range: 0.011-0.63 *μ*g L^-1^, Fig. S3). Therefore, these samples represent the ocean environments in which small phytoplankton are most important, and our isolates were matched to metabarcode-based operational taxonomic units (OTUs) (Materials and Methods). On average the 13 OTUs studied here account for 31% of all metabarcode reads from non-dinoflagellate phytoplankton in this size fraction (Materials and Methods).

All mixotroph OTUs except one (*Florenciella* sp.) had greater relative abundance in surface samples than deep chlorophyll maximum (DCM) samples, while all autotroph OTUs had greater relative abundance in DCM samples (Fig. S4). Furthermore, the surface:DCM relative abundance ratio is correlated with grazing ability (i.e., specific clearance rate when fed *Prochlorococcus*), such that better grazers have shallower distributions (Fig. 2a). The statistical relationship between grazing ability and depth niche is clearest when autotrophs and mixotrophs are both included - the 95% credible interval for the effect of clearance rate on the depth ratio is [0.18, 0.71], and *R*^2^ = 0.55 for the relationship between these two variables. However, the trend remains when only mixotrophs are considered (95% CI = [-0.03, 0.72], R^2^ = 0.38), indicating that depth differences among mixotrophs correlate with their relative grazing abilities.

**Figure 2.**
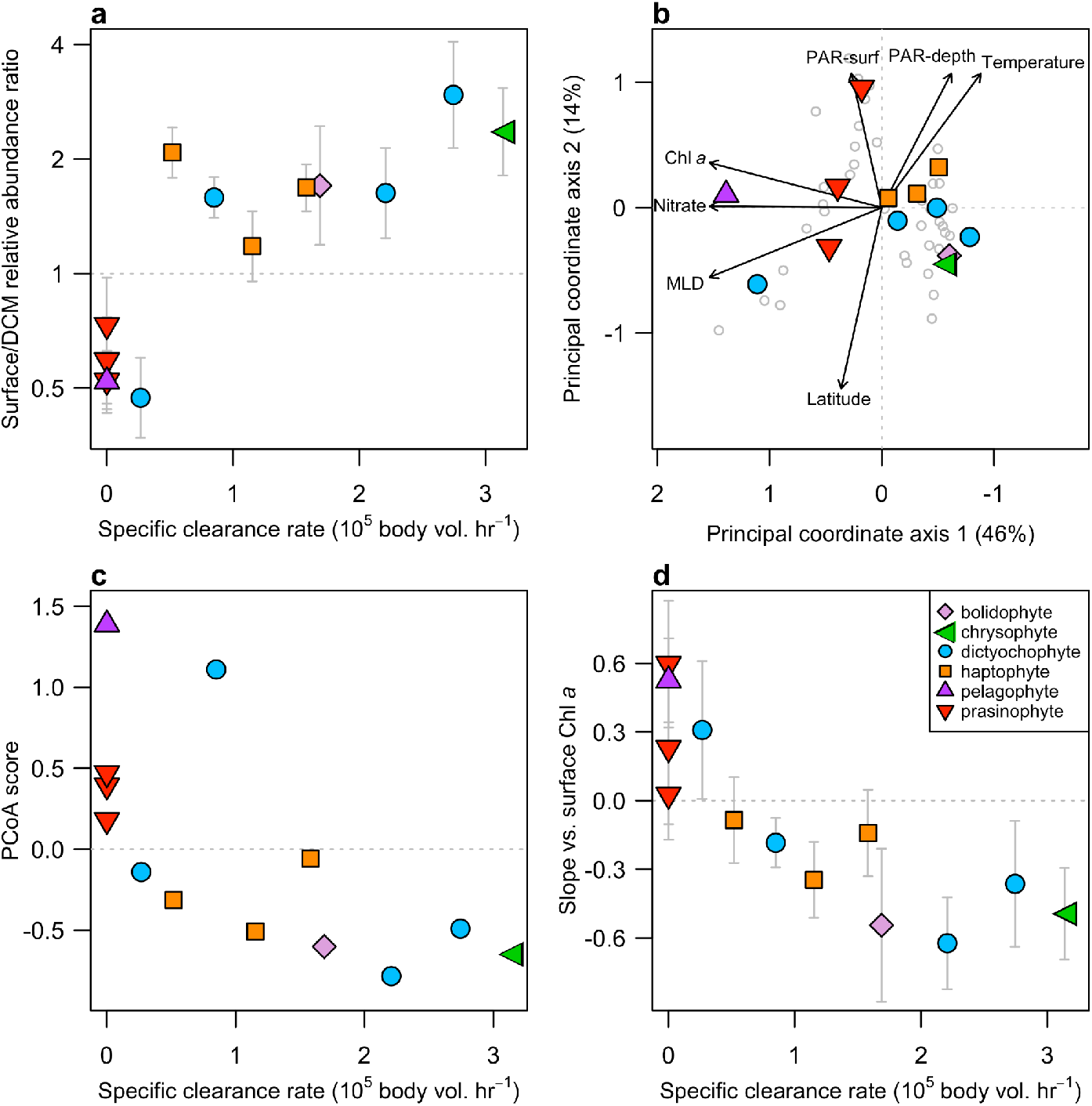
Environmental niches of 13 phytoplankton OTUs compared to grazing ability of corresponding isolates. (**A**) Y-axis is the ratio of OTU relative abundance in surface vs. deep chlorophyll maximum (DCM) samples, derived from a GLMM fit to *Tara* Oceans metabarcodes in the 0.8-5 *μ*m size fraction. X-axis is grazing ability, quantified as specific clearance rate when fed *Prochlorococcus,* and strains exhibiting no ability to grow when fed prey are given zeros. (**B**) The first two axes from a principal coordinate analysis (PCoA) of OTU composition across surface samples. Vectors represent the correlations between environmental variables and the axes. PAR-surf = photosynthetically active radiation at the surface, PAR-depth = PAR at the sample depth, Latitude = absolute latitude. (**C**) OTU position along the first PCoA axis compared to grazing ability. (**D**) Y-axis is the slope of OTU relative abundance vs. chlorophyll *a* concentration in surface samples, derived from a GLMM fit to *Tara* Oceans metabarcodes in the 0.8-5 *μ*m size fraction. X-axis is grazing ability as described for (**A**).

Much of the variation in OTU composition across surface samples can be explained by a single principle coordinate axis (46%; Fig. 2b). This axis is strongly positively correlated with Chl *a*, nitrate, and mixed layer depth, and moderately negatively correlated with temperature and photosynthetically active radiation (PAR) at the sample depth (Fig. 2b). Therefore, this axis likely represents community structure driven by stratification, with less stratified waters having deeper mixed layers, greater nutrient supply and Chl *a*, and PAR diminished by greater pigment concentration. The position of OTUs along this axis is correlated with grazing ability (r = −0.67, p = 0.011; Fig. 2c), with autotrophs and slower-grazing mixotrophs more abundant under less stratified conditions. There is also a nonsignificant trend when only considering the nine mixotrophs (r = −0.5, p = 0.17). A similar but stronger pattern is found when considering niche differences across Chl *a* gradients (Fig. S5). The four autotrophs and one mixotroph (*Florenciella* sp.) increase in relative abundance as Chl *a* increases, while the other mixotrophs decline, and better grazers show a steeper decline with increasing Chl *a* (Fig. 2d). The relationship between grazing ability and Chl *a* niche is clearest when autotrophs and mixotrophs are both included (95% CI = [−0.39, −0.14], R^2^ = 0.9), but remains when only mixotrophs are considered (95% CI = [−0.37, −0.06], R^2^ = 0.8). Increasing the phylogenetic scale of these analyses, such that grazing abilities of isolates are matched to average niches of their respective families or orders, yield similar patterns, implying that the trait-niche relationships of OTUs are reflective of broader phylogenetic structure in these communities (Fig. S6).

The trait-niche relationships in Fig. 2 indicate that there are parallel changes in community structure when transitioning from deeper to shallower depths in the euphotic zone, and when transitioning from less stratified, high Chl *a* locations to more stratified, lower Chl *a* locations. Both of these gradients are associated with shifts from autotrophs and slower-grazing mixotrophs to faster grazing mixotrophs (Fig. 3); they are also associated with concomitant changes in the availability of nutrients and light, resources known to affect the relative fitness of different trophic strategies. To consider the distinct roles of nutrient and light gradients, we utilize a trait-based model.

**Figure 3.**
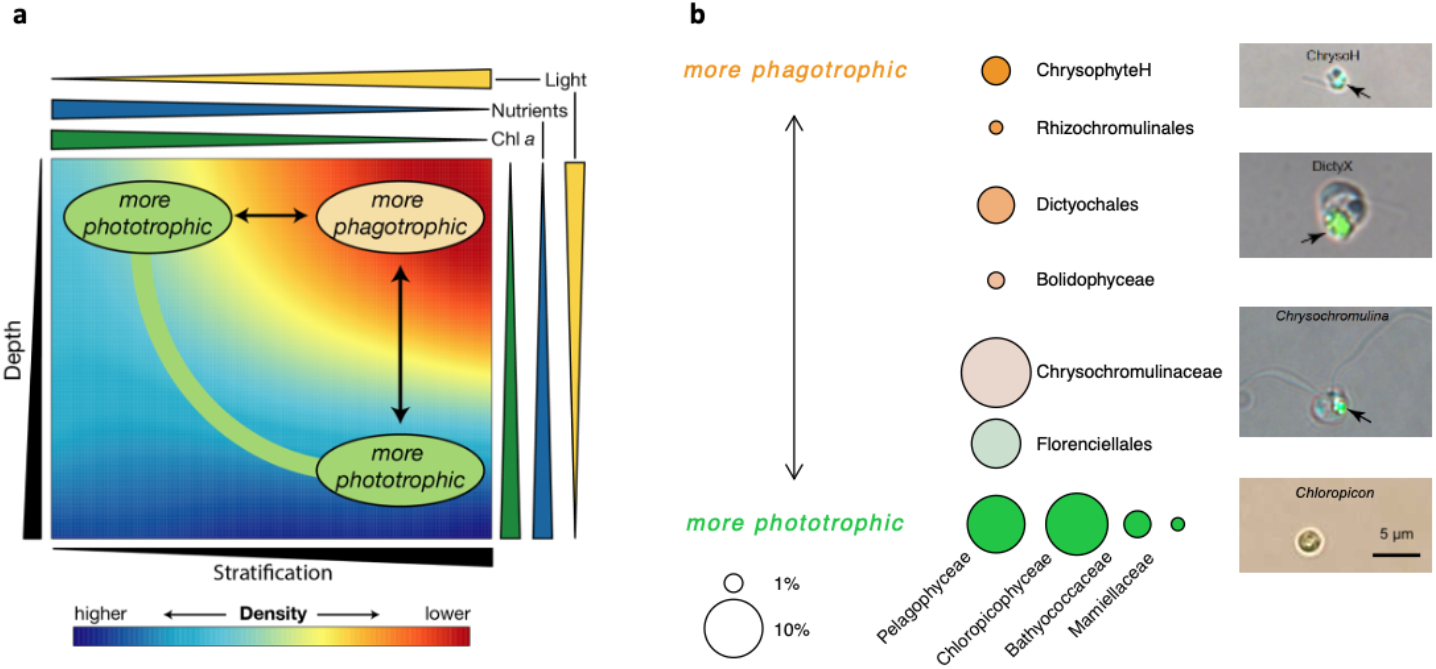
(**A**) Diagram of how depth and stratification gradients lead to parallel changes in phytoplankton community trait structure. Eukaryotic phytoplankton become more phagotrophic at shallower depths and in more stratified surface waters. Greater stratification decreases nutrient supply across the nutricline, and increases light intensity by reducing mixed layer depth and light-absorbing phytoplankton pigments. Moving from deeper to shallower depths is also associated with less nutrients and more light, and lower Chl *a*. (**B**) Diagram of the hypothesized spectrum of phytoplankton trophic strategies, based on traits of isolates and niches of OTUs and broader taxa (Figs. 1, 2, S6). Bubble area is mean relative abundance in surface *Tara* Oceans samples, within the 0.8-5 *μ*m size fraction of phytoplankton (as defined in Materials & Methods). Images show example isolates from common taxa, arrows point to fluorescent beads ingested by mixotrophs as described in Li et al. (2022). In panel (**A**), moving across depth or stratification gradients leads to parallel shifts in relative abundance of taxa across the spectrum in panel (**B**).

### Trait-based model of trophic strategies across environmental gradients

We analyzed a trait-based model where a spectrum of populations with different traits compete for dissolved nutrients and bacterial prey at a defined irradiance. Photosynthesis and phagocytosis were parameterized to be consistent with experimental data, which required a relatively strong tradeoff. We assume the ability to ingest prey causes a steep decline in photosynthetic capacity, as well as an increase in respiratory costs (Fig. S7). The model predicts that an increase in nutrient supply causes autotrophs to increase relative to mixotrophs, and at the same time mixotrophic strategies that invest more in phototrophy increase relative to strategies that invest less in phototrophy (i.e., the mean trophic strategy parameter declines; Fig. 4). Across a gradient of irradiance, autotrophs become relatively more abundant at low irradiances, and mixotrophs become more phototrophic as irradiance declines (Fig. 4). The effect of irradiance is sensitive to the tradeoff assumptions - if photosynthetic capacity of phagotrophs is weakly penalized, then mixotrophs can outcompete heterotrophs and autotrophs at lower irradiances, and mixotrophs become more phagotrophic under those conditions (Fig. S8). In contrast, if the greater respiratory cost of phagotrophy is removed, then this benefits heterotrophs, who strongly suppress mixotrophs at lower irradiances, which indirectly allows autotrophs to prosper under low light (Fig. S9). In contrast to the irradiance gradient, the effect of the nutrient gradient is not qualitatively affected by tradeoff assumptions.

**Figure 4.**
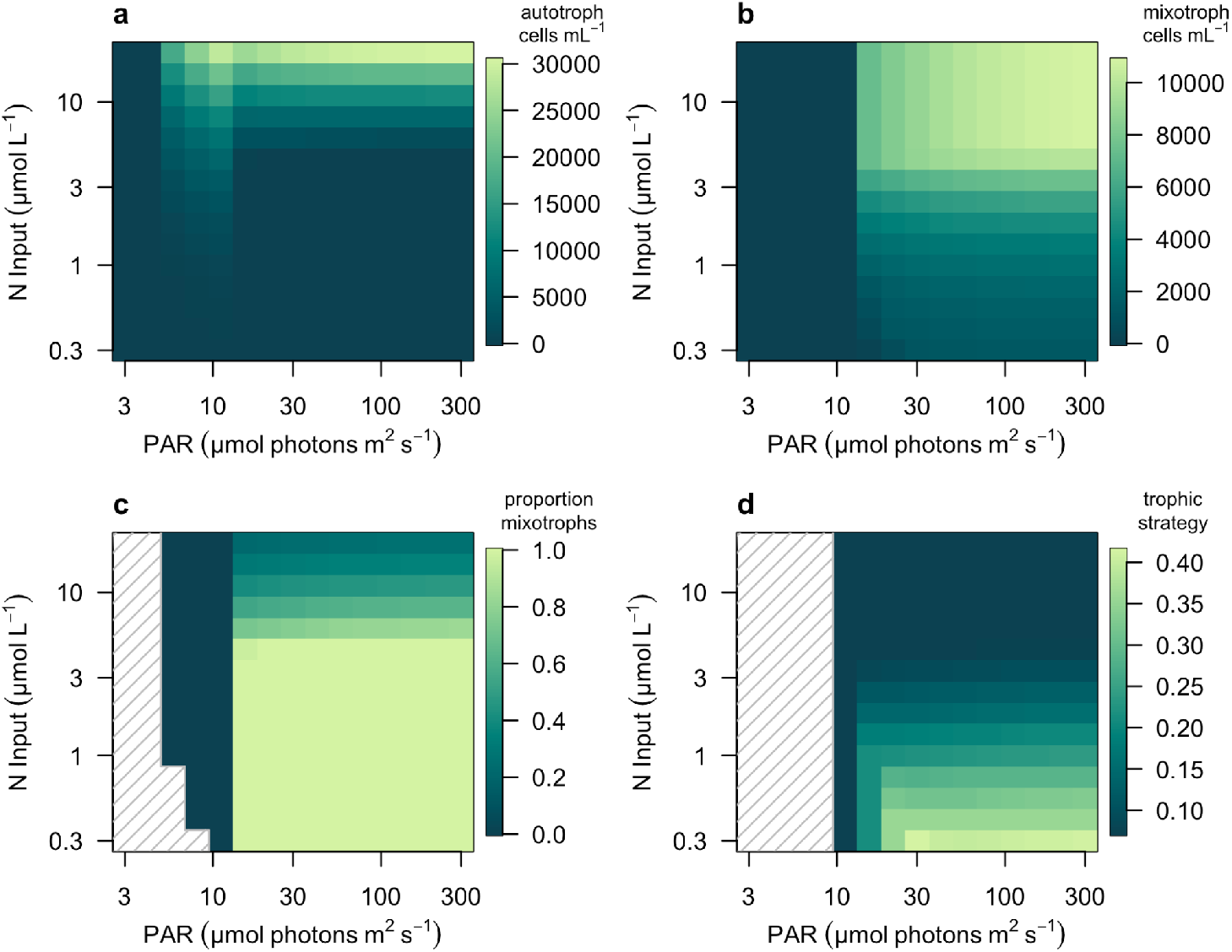
Modeled trophic strategies vs. gradients of nitrogen input and irradiance. (**A**) Concentration of autotrophs. (**B**) Concentration of mixotrophs. (**C**) Mixotrophs as a proportion of all phytoplankton. (**D**) Trophic strategy of the persisting mixotroph population, where strategy is the parameter *x* (see Materials and Methods) which ranges from 0 (autotroph) to 1 (heterotroph). Regions with cross-hatching denote where phytoplankton did not persist (panel **C**) or mixotrophs did not persist (panel **D**). For brevity the heterotroph population is omitted (see Figs. S8-9).

In total the model results suggest that strong relationships between grazing ability and environmental niches across multiple gradients (Figs. 2-3) may be driven by nutrient supply and potentially irradiance. Moving from shallow to deep within the euphotic zone is associated with increasing nutrient supply as well as declining irradiance. Likewise, nutrient supply increases and irradiance declines when moving from more stratified / low Chl *a* waters to less stratified / high Chl *a* waters. Nutrient supply across these gradients should favor relatively phagotrophic phytoplankton at shallower depths and in more stratified water columns (Fig. 4). The role of irradiance is more complex, but under relatively strong tradeoffs consistent with our data the greater irradiance at shallower depths and in more stratified water columns should also favor more phagotrophic phytoplankton (Fig. 4).

## Conclusions

Our results document tradeoffs among diverse autotrophic and mixotrophic phytoplankton, and demonstrate predictable shifts in their trophic strategies across ocean environments. It is noteworthy that a single axis of phototrophy vs. phagotrophy seems to capture much of the ecological variation among taxa from many deeply branching clades. This suggests that fundamental constraints on physiology may lead to ‘universal’ tradeoffs that underlie trait diversity, community structure, and ecosystem function. The approach we have taken can only characterize the subset of the community that we have isolated. We have considerably expanded the range of oceanic isolates available for experimentation with this work, but a fuller accounting of eukaryotic diversity will require isolation of additional common clades, as well as *in situ* methods for quantifying trait variation. In addition, the relative fitness of mixotrophic strategies depends on competition with heterotrophs, which have not been examined in this study. Isolation of prevalent open-ocean heterotrophs, and comparison of their traits and niches to co-occurring mixotrophs, is an important direction for future research.

The observed patterns in community trait structure imply gradients in ecosystem functioning, such that phagotrophically supported primary production may be greatest at shallow depths and in highly stratified waters, which could alter trophic transfer efficiency and nutrient cycling (*20, 21*). There are many potential drivers of stratification across the *Tara* Oceans survey locations, and one possibility is seasonal cycles that were at different stages at the different sampling stations. A seasonal component to the niche differences in Fig. 2b-d would mean that seasonal patterns among small eukaryotes are similar to those seen in larger microphytoplankton, where mixotrophic dinoflagellates become more abundant than autotrophic diatoms during stratified summer conditions (*33*), consistent with an optimality-based model of succession among trophic strategies (*24*). Our results also suggest that climate change, which is increasing ocean stratification (*34*), is also making phytoplankton communities more phagotrophic.

A previous analysis of nanoplankton found that mixotrophs increase relative to autotrophs at lower latitudes, but both mixotrophs and autotrophs increase relative to heterotrophs in productive coastal waters (*29*). The latter pattern appears to contradict the current results, but it could be consistent if the increase in mixotroph abundance in coastal waters was driven by the relatively autotrophic dictyochophyte *Florenciella*. This genus is globally abundant (*17*), grazes relatively slowly (*17*) (Fig. 2), grows quickly on dissolved nutrients (*35*) (Fig. 1), and contains the one mixotroph OTU in our analysis that increases in relative abundance across a Chl *a* gradient (or increases at the DCM; Fig. 2; Figs. S4-5). In general, the wide variation in traits among mixotrophs, which is correlated with environmental gradients, implies that bulk measurements of mixotroph abundance may obscure significant shifts in community function.

Recent work has shown that mixotrophs that do not make their own chloroplasts have biogeographies that depend on the mode of chloroplast acquisition (*36*), and modeling shows that non-constitutive mixotrophs may have a different niche than the constitutive form (*37*). Integrating the spectrum of constitutive and acquired mixotrophies into a unified analysis of competitive outcomes should provide a more complete understanding of trophic strategies among unicellular protists. Finally, it is noteworthy that the correlation of grazing ability with a population’s Chl *a* niche (Fig. 2d) provides a link to remote sensing. Phytoplankton community trophic strategy may be predictable at a global scale using remotely sensed Chl *a*, and this may also provide a route for phagotrophy to be better incorporated into models of primary production.

## Materials and Methods

### Strain isolation and cultivation

Isolation methods for most strains used in this study were described in Li et al. (*17*). Briefly, the majority were isolated from the euphotic zone at Station ALOHA (22°45’ N, 158°00’ W) in 2019, enriched using Keller (K) medium with a 20-fold reduction in mineral nutrients and addition of *Prochlorococcus* (MIT9301) at ~5×10^6^ cells mL^-1^. Five strains were isolated from previous samples at the same location, enriched with full K medium or K medium without added nitrogen. Four strains used in the present study (*Ostreococcus, Chloropicon, Micromonas, Pelagomonas*) were not described in Li et al. (*17*). These strains were isolated from the same location, enriched using full K medium. All strains were rendered unialgal but not axenic, maintained at 24°C in 0.2 *μ*m-filtered and autoclaved ALOHA seawater, under a 12:12 light:dark cycle with irradiance ~70 *μ*M photons m^-2^ s^-1^. Mixotrophs (as described in Li et al. (*17*)) were maintained in K medium without added nitrogen, amended with *Prochlorococcus* prey. The four strains not previously described were maintained in full K medium. Strain taxonomy was characterized with phylogenetic analysis of 18S rDNA as described in Li et al. (*17*).

### Mixotrophy assays

Eleven strains were used to compare phototrophic growth abilities with the ability to grow when fed *Prochlorococcus*. Seven strains were previously shown to consume *Prochlorococcus* and grow when fed *Prochlorococcus* as the only added nitrogen source: a chrysophyte from environmental clade H (hereafter ChrysoH), a bolidophyte in the genus *Triparma*, two haptophytes in the genus *Chrysochromulina*, two dictyochophytes in the genus *Florenciella*, and one undescribed chlorarachniophyte (hereafter ChloraX) (*17*). Four strains were newly assayed for ability to grow when fed *Prochlorococcus*: three prasinophytes from the genera *Ostreococcus, Chloropicon*, and *Micromonas*, and one pelagophyte in the genus *Pelagomonas*. Three previously assayed mixotrophs were included in the new assays (two *Chrysochromulina* and ChrysoH) as positive controls. Strains were inoculated into K medium without added nitrogen at ~10^3^ cells mL^-1^ and monitored for eight days to allow consumption of residual nitrogen, at which point *Prochlorococcus* was added at ~10^6^ cells mL^-1^. Cultures were monitored for eight more days for evidence of growth and compared to control cultures without added *Prochlorococcus*.

### Phototrophic growth measurements

The eleven strains used in mixotrophy assays were also tested for phototrophic growth ability, i.e., the ability to grow using light and dissolved nutrients as resources. Cultures were inoculated at ~10^2^-10^3^ cells mL^-1^ into tissue culture flasks containing 20 mL K medium, at two irradiances (10 and 100 *μ*mol photons m^-2^ s^-1^), referred to as ‘low’ and ‘high’ light, respectively. Samples were taken every 1-3 days and incubated in a final volume of 0.5% glutaraldehyde for 15 minutes before flash freezing in liquid nitrogen and storage at −80°C, followed by counts with flow cytometry. All strains were acclimated by passaging through at least one batch culture in the experimental conditions, before collecting data to estimate growth rates. Some strains grew at high irradiance but consistently failed to grow at low irradiance after repeated inoculations, as noted in the main text. One strain (ChrysoH) was unable to grow phototrophically (i.e., without added prey), although it grew readily with added prey. For strains that grew, growth rate was estimated using at least two replicates in all cases, except one *Chrysochromulina* strain for which one high light growth rate was obtained.

Growth rates were estimated by fitting nonlinear growth models to cell concentrations over time. For cultures that exhibited a lag before exponential growth, a growth model with a lag phase and carrying capacity was fit: 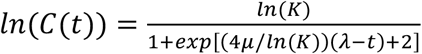, where *C*(*t*) is cell concentration at time *t, μ* is the exponential growth rate, *K* is carrying capacity (stationary density), and *λ* is the inflection point where growth rate equals *μ* (*38*). For cultures that exhibited no lag, a logistic growth model was fit: 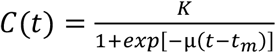, where *t*_m_ is the inflection point of the logistic curve and *C*(*t*), *K*, and *μ* have the same meaning. The models were fit by maximum likelihood with the R package bbmle (*39*). In cases where cell concentration declined after reaching maximal abundance, the decline phase was omitted to allow the model to fit to the sigmoidal portion of the growth curve.

### *Tara* Oceans OTUs

In order to compare traits of our isolates to the environmental niches of their populations, or closely related populations, we utilized the *Tara* Oceans eukaryotic plankton diversity dataset (*40*). This dataset contains size-fractionated 18S-V9 rDNA metabarcodes from 40 stations in the sunlit ocean (http://taraoceans.sb-roscoff.fr/EukDiv/; Fig. S3). Using 27 strains for which we previously measured clearance rates when consuming *Prochlorococcus* (*17*), plus four additional strains assayed in this study (*Ostreococcus, Chloropicon, Micromonas, Pelagomonas*), we matched strains to OTUs from the *Tara* Oceans dataset (Table S2). Near-full length 18S rDNA sequences of our isolates were compared to all OTU 18S-V9 rDNA reference sequences using nucleotide BLAST. In nearly all cases the OTU with the lowest E-score was frequent enough to analyze abundance patterns across samples (i.e., thousands of reads or more), and this OTU was chosen for further analysis. In some cases there was a second OTU with an equivalent E-score but <10 reads, and this OTU was not used. In one case (DictyX) the OTUs with the two lowest E-scores had <10 reads, and the third ranking OTU was chosen. In one case (*Chloropicon*) two abundant OTUs had the same E-score, and their reads were summed in each sample for further analysis (choosing one OTU produced similar results). The chlorarachniophyte strain ChloraX was not similar to any OTU abundant enough for further analysis. *Florenciella* strains were divided into two groups, one group that best matched a *Florenciella parvula* OTU and one that matched an OTU from an undescribed *Florenciella* species. In all cases taxonomic annotation of the *Tara* Oceans OTUs was consistent with isolate taxonomy independently derived by phylogenetic analysis of isolate 18S rDNA and related sequences from GenBank and the PR^2^ database (*41*).

### Statistical analyses

We asked whether the environmental niches of phytoplankton OTUs are correlated with their grazing ability. Grazing ability was quantified as body volume-specific clearance rate when fed ~10^6^ cells mL^-1^ *Prochlorococcus* (*17, 41*), and this trait was used because we have measured it on a large number of isolates. As described previously, the clearance rates measured with these isolates approximate their maximum clearance rates, because prey concentrations were low enough to not saturate the ingestion rate (*17*). Isolates determined to be autotrophic by our mixotrophy assays were given a grazing ability of zero. This yielded a total of 13 OTUs for which niches could be compared to grazing ability. Metabarcodes from the ‘pico/nano’ 0.8-5 *μ*m size fraction were used, as all of our isolates are within this size class.

We took two approaches to test whether grazing ability correlates with niche differences. We used principal coordinate analysis to ordinate major axes of compositional variation among our focal OTUs. To interpret drivers of composition we then correlated the first principal coordinate axis with environmental variables: Chl *a* concentration (HPLC), photosynthetically active radiation (PAR) at the sea surface, PAR at the sample depth, nitrate concentration, sea surface temperature, mixed layer depth, and absolute latitude. All variables were taken from ancillary *Tara* Oceans datasets (*42–44*). Finally, the position of OTUs along the first axis was compared to their grazing ability.

We also asked whether grazing ability was correlated with OTU responses to specific environmental variables chosen *a priori:* depth (surface [3-7 m] vs. deep chlorophyll maximum [DCM]), and Chl *a* concentration. When using Chl *a* as the predictor only surface samples were used, and one station was withheld because it had much higher Chl *a* concentration (5.5 *μ*g L^-1^) than the other stations (0.011-0.63 *μ*g L^-1^). Samples without Chl *a* data were excluded in this analysis. For depth and Chl *a* generalized linear mixed models (GLMMs) were fit with OTU relative abundances modeled using the beta-binomial distribution with a logit link function: logit(*p*_ij_) = Int_i_ + Sample_j_ + (CR_i_*CReff + slope_i_)*Env_j_, reads_ij_ ~ BetaBinom(*p*_ij_, *V*_i_, *N*_j_) Here *p*_ij_ is the probability that a metabarcode read in sample *j* is from OTU *i*, Inti is an OTU-specific random intercept capturing variation in mean relative abundance across OTUs, Samplej is a random effect capturing variation in mean relative abundance of all OTUs across samples, CR is specific clearance rate of OTU *i*, CReff is the effect of clearance rate on OTU responses to the environment, slope_i_ is a species-specific random slope capturing variation in environmental responses not attributable to CR, Env_j_ is the value of the environmental variable in sample *j*, reads_ij_ is number of reads of OTU *i* in sample *j, V*_i_ is an OTU-specific dispersion parameter, and *N*_j_ is the total number of phytoplankton reads in sample *j*. In summary, this model quantifies whether the relationship between relative abundance and an environmental variable for an OTU is predicted by that OTU’s clearance rate. The GLMM approach is appropriate because it models # of reads while accounting for variation in total reads, and allows for uncertainty in relative abundances and environmental relationships while quantifying CReff (*45, 46*). The assumption of logit-linear environmental responses was appropriate based on visual inspection of the data (Fig. S5). To account for potential phylogenetic correlation in OTU environmental responses we also fit models with additional random effects for taxon (haptophyte / dictyochophyte / prasinophyte / chrysophyte / pelagophyte / bolidophyte), but in all cases variance of these effects was zero. Models were fit in R using the package brms, which implements bayesian regression models via the software Stan (*47*).

To define the total number of phytoplankton reads we summed the reads of known phytoplankton taxa (*Tara* Oceans ‘taxogroups’: Bacillariophyta, Bolidophyceae, Chlorarachnea, Chlorophyceae, Chrysophycea/Synurophyceae, Cryptophyta, Dictyochophyceae, Euglenida, Glaucocystophyta, Haptophyta, Mamiellophyceae, Other Archaeplastida, Other Chlorophyta, Pelagophyceae, Phaeophyceae, Pinguiophyceae, Prasino-Clade-7, Pyramimonadales, Raphidophyceae, Rhodophyta, Trebouxiophyceae). Dinoflagellates were excluded because of the difficulty in assigning phototrophic vs. heterotrophic status to all taxa, and because nearly all dinoflagellate reads were from a single, poorly annotated OTU that was also highly abundant in larger size fractions. We also excluded a small number of taxa within the taxogroups listed above that are known to be heterotrophic. However, neither the exclusion of dinoflagellates nor the heterotrophs within majority-phytoplankton taxogroups qualitatively changes our results.

### Trait-based model of trophic strategy competition

In this study we perform new analyses of a model similar to that described by Edwards (*29*). The model describes population growth of single-celled protists where the potential limiting factors are dissolved nutrients, bacterial prey, and light. The dynamic variables are the abundance of protist population *i* (*P*_i_) which could be mixotrophic or heterotrophic or autotrophic, the internal nutrient quota of protist *i* (*Q*_i_) the abundance of bacterial prey (*B*), and the concentration of dissolved nutrient (*N*):

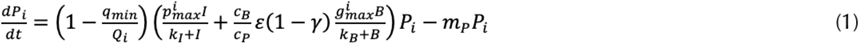

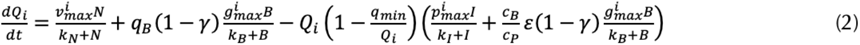

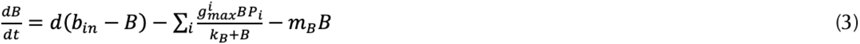

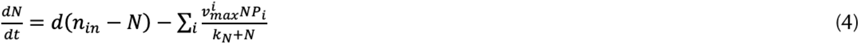

In equation 1, the term 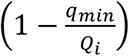 describes limitation of growth by the internal nutrient quota *Q*_i_, with minimum subsistence quota *q*_min_. This is multiplied by a term describing limitation by the energy and organic carbon obtained from photosynthesis and/or prey ingestion, which has two components: (1) photosynthesis, 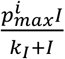, where *p*^i^_max_ is the maximum net photosynthetic rate for protist *i, k*_i_ is the half saturation constant, and *I* is irradiance; (2) ingestion, 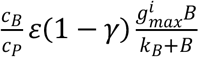, where *c*_B_ is bacterial carbon per cell, *c*_P_ is protist carbon per cell, *ε* is the net growth efficiency, *γ* is the fraction of prey unconsumed or egested, *g*^i^_max_ is the maximum ingestion rate, and *k*_B_ is the half saturation constant for ingestion. This growth model treats nutrient and energy/carbon limitation as multiplicative, consistent with the fact that autotrophic growth is sensitive to irradiance under nutrient limitation, and vice versa (*48–50*); the same is true for heterotrophic bacteria consuming organic carbon and inorganic nitrogen (*51*). The protist population declines due to a constant per capita mortality rate *m*_P_.

Equation 2 follows the internal nutrient quota, which increases due to (1) dissolved nutrient uptake at rate 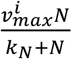, where 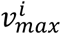 is the maximum uptake rate and *k*_N_ is the halfsaturation constant; and (2) prey ingestion at rate 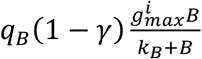, where *q*_B_ is the bacterial nutrient per cell. The nutrient quota declines due to carbon acquisition during growth, which is the third term in the equation. Eqns. (1) and (2) can represent mixotrophic growth, pure autotrophy (by setting *g*^i^_max_ to 0), or pure heterotrophy (by setting 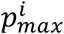 to 0).

Equation 3 follows bacterial abundance dynamics, which could include autotrophic and heterotrophic bacterial prey. Bacterial growth is not modeled explicitly because the primary focus is how different protist trophic strategies respond to the supply of nutrients, bacterial prey, and light. Therefore, the supply of bacteria is controlled by mixing bacteria at concentration *b*_in_ into the system at rate *d* (first term in eqn. 3). The second term in eqn. 3 is loss due to combined consumption by all grazers, and the third term is a background constant mortality at per capita rate *m*_B_. Equation 4 follows the dissolved nutrient, which is parameterized to represent nitrogen (forms of nitrogen are not distinguished), but could generically represent any limiting nutrient. The first term is the supply of nutrient of concentration *n*_in_ at rate *d*, and the second term is the uptake of nutrient by the flagellate populations.

We assume that species can vary continuously in allocating resources to phototrophy or phagotrophy. This tradeoff affects the rate of photosynthesis at all irradiances through the parameter 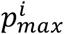, and the rate of prey ingestion at all prey concentrations through the parameter 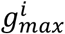. For simplicity we assume that ability to take up and assimilate dissolved nutrients is proportional to 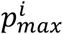, yielding a single tradeoff axis of phototrophy vs. phagotrophy. If position on the tradeoff axis is *x*, then 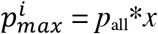 and 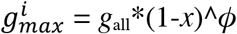, where *p*_all_ and *g*_all_ are the maximal rates obtained by autotrophs and heterotrophs, respectively. The parameter ϕ controls the curvature of the tradeoff and therefore its strength: *ϕ* > 1 penalizes mixotrophs for a generalized strategy, while *ϕ* < 1 rewards them (*29*). Motivated by our experimental results, we also consider that phytoplankton engaged in phagotrophy may have greater respiratory demand, resulting in a steep decline in phototrophic growth at low irradiances. This is implemented by making the mortality/loss constant *m*_P_, which represents maintenance respiration as well as other losses, a function of trophic strategy: *m*_P_ = *m*_0_ + *m*_1_*(1-*x*). In our simulations we use several tradeoff scenarios. The baseline scenario assumes the phototrophy-phagotrophy tradeoff is strong (*ϕ* = 1.5) and *m*_P_ increases with phagotrophy, because we found these assumptions were necessary to achieve growth-irradiance relationships consistent with our experimental data (Fig. 1, Fig. S8). This scenario is compared to outcomes when the tradeoff is weak (*ϕ* = 0.8) and/or *m*_P_ does not increase with phagotrophy. Units and values used for all model parameters are given in Table S3.

To ask how community structure varies across environmental gradients we initialized communities with 20 species at low density (10 cells mL^-1^) ranging from *x* = 0 to *x* = 1, and solved the dynamics numerically until a steady state attractor was reached. Nutrient supply (parameterized to represent nitrogen, though the results can represent any limiting nutrient) was manipulated by varying the input concentration *N*_in_ from 0.3 to 20 *μ*mol L^-1^. Irradiance was varied from 3 to 300 *μ*mol photons m^-2^ s^-1^, with 20*20 nutrient and light combinations. In all cases we found that the community reached a steady state attractor which was uninvasible by any trophic strategy that had gone extinct during community assembly. In other words, the community structure reported for a set of parameters is an evolutionarily stable community (*52*).

## Acknowledgments

We thank the personnel in the Hawai’i Ocean Time-series program (NSF award 12-60164) for assistance with collecting water samples and Karen Selph for assistance with flow cytometry.

## Supplementary Figures

**Figure S1.**
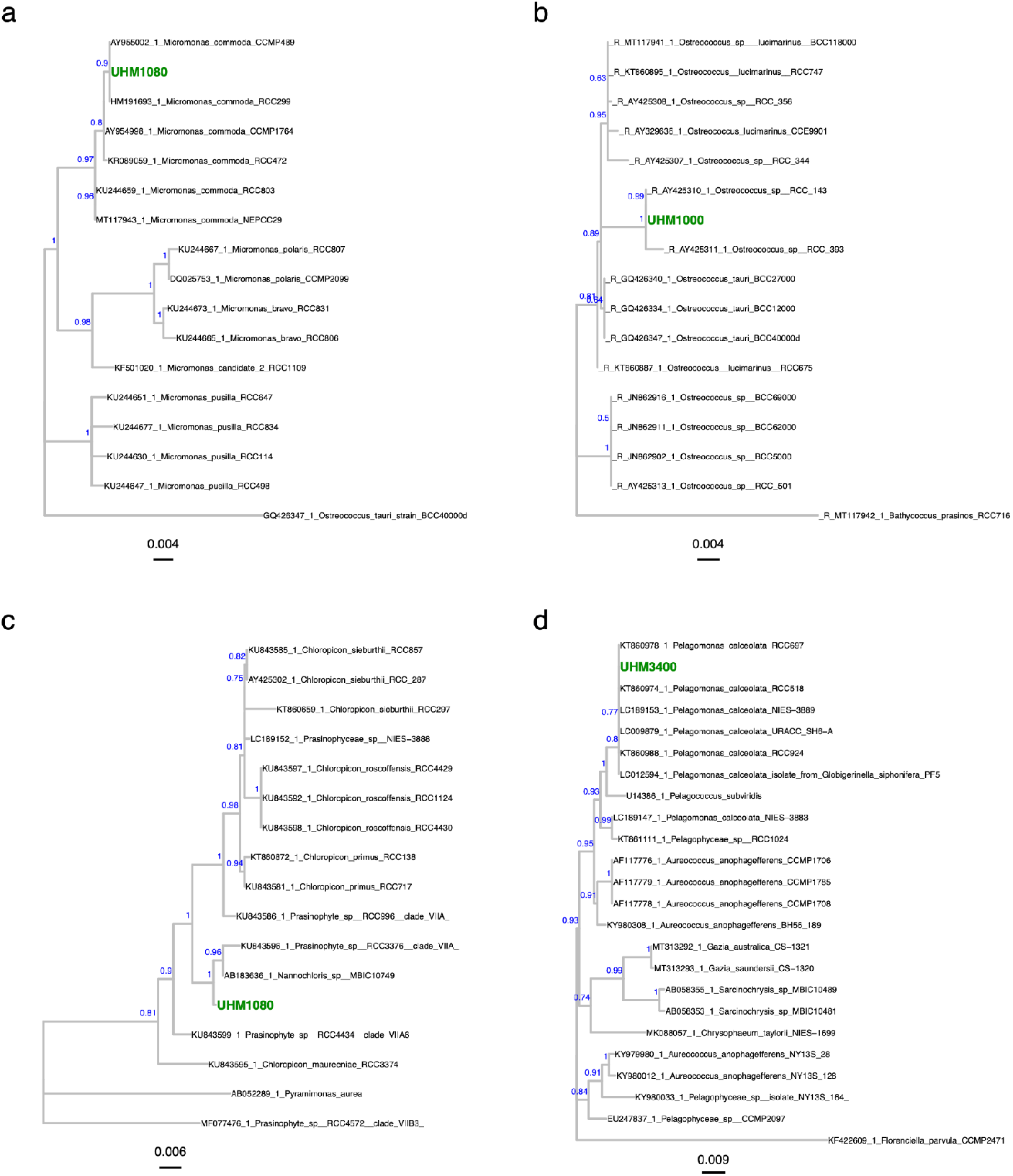
Phylogenetic placement of 18S rDNA genes of four previously undescribed isolates, putatively assigned to the genera *Micromonas* (a), *Pelagomonas* (b), *Chloropicon* (c), and *Ostreococcus* (d). For each tree a selection of closely related 18S rDNA sequences from cultured isolates were retrieved from GenBank and aligned with MAFFT. Trees were fit with raxmlGUI 2.0, numbers on nodes are bootstrap support (out of 1000) and scale bars are nucleotide substitutions per site. The new isolates are listed in bold green text.

**Figure S2.**
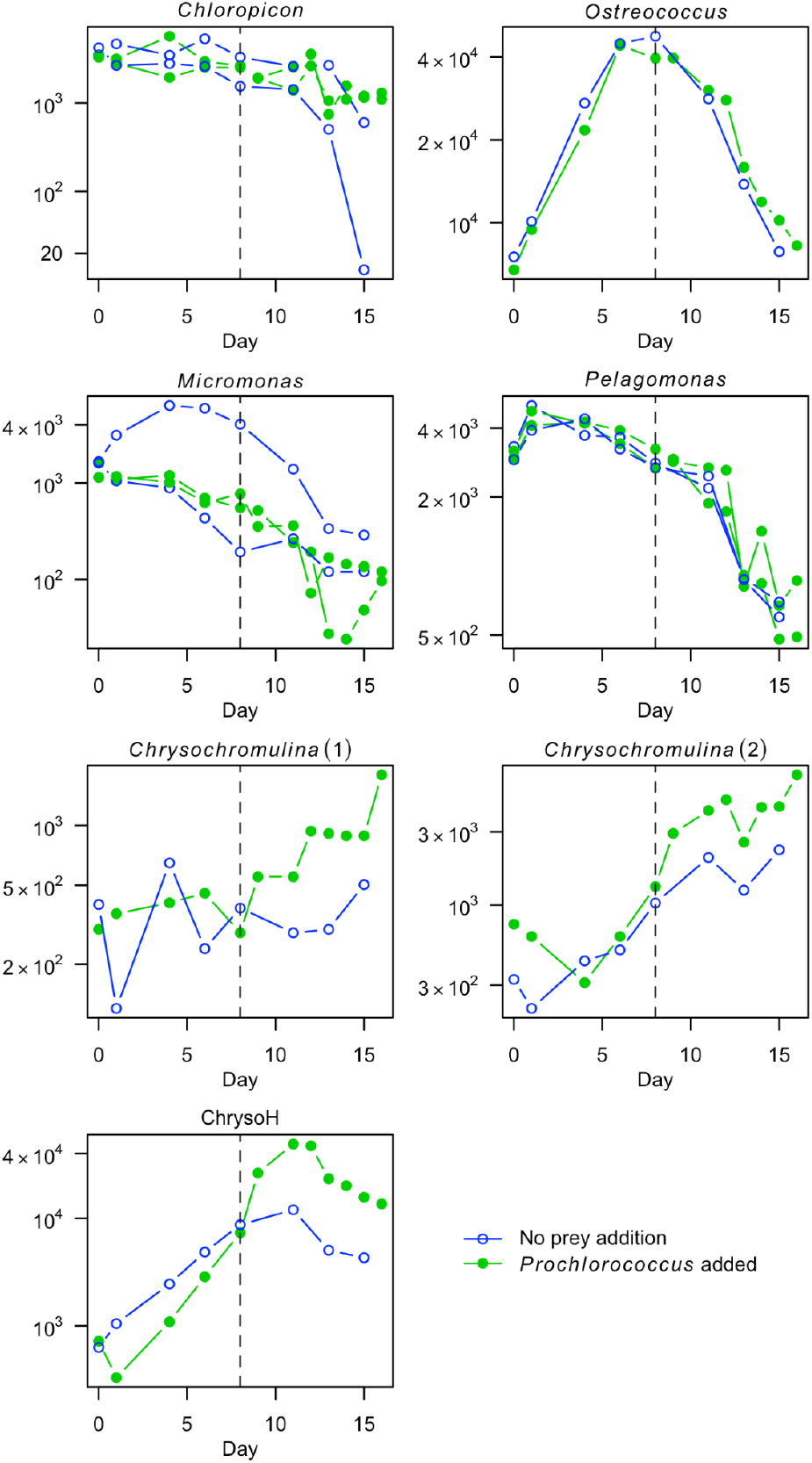
Assay of mixotrophic growth ability for seven phytoplankton strains. Strains were inoculated into K medium without added nitrogen, and after 8 days (dashed line) *Prochlorococcus* were added at ~10^6^ cells mL^-1^. The strains previously shown to grow when fed *Prochlorococcus (Chrysochromulina* (1), *Chrysochromulina* (2), and ChrysoH) increased relative to control cultures (no prey added), while the remaining strains (*Ostreococcus, Chloropicon, Micromonas, Pelagomonas*) did not. All additional strains used in this study were previously shown to consume *Prochlorococcus* and grow mixotrophically (*17*).

**Figure S3.**
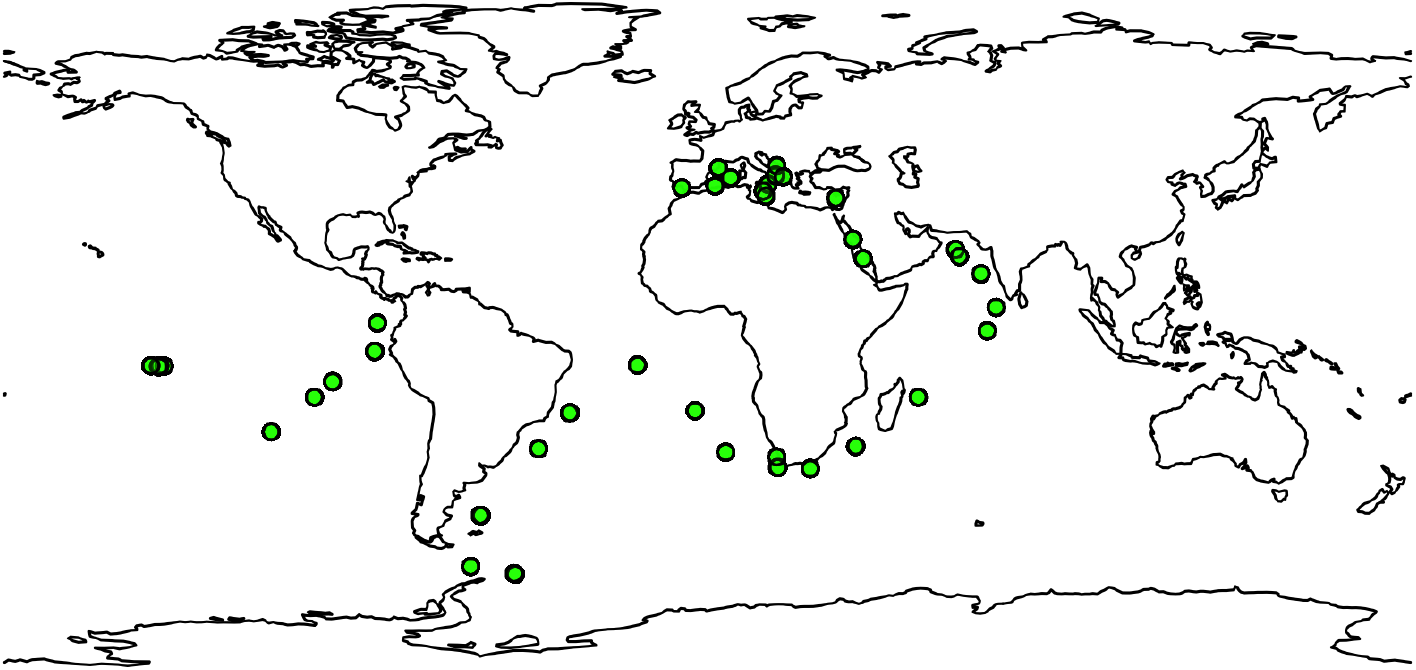
Locations of the *Tara* Oceans stations at which 0.8-5 *μ*m size fraction 18S V9 rDNA metabarcodes were sampled (*40*).

**Figure S4.**
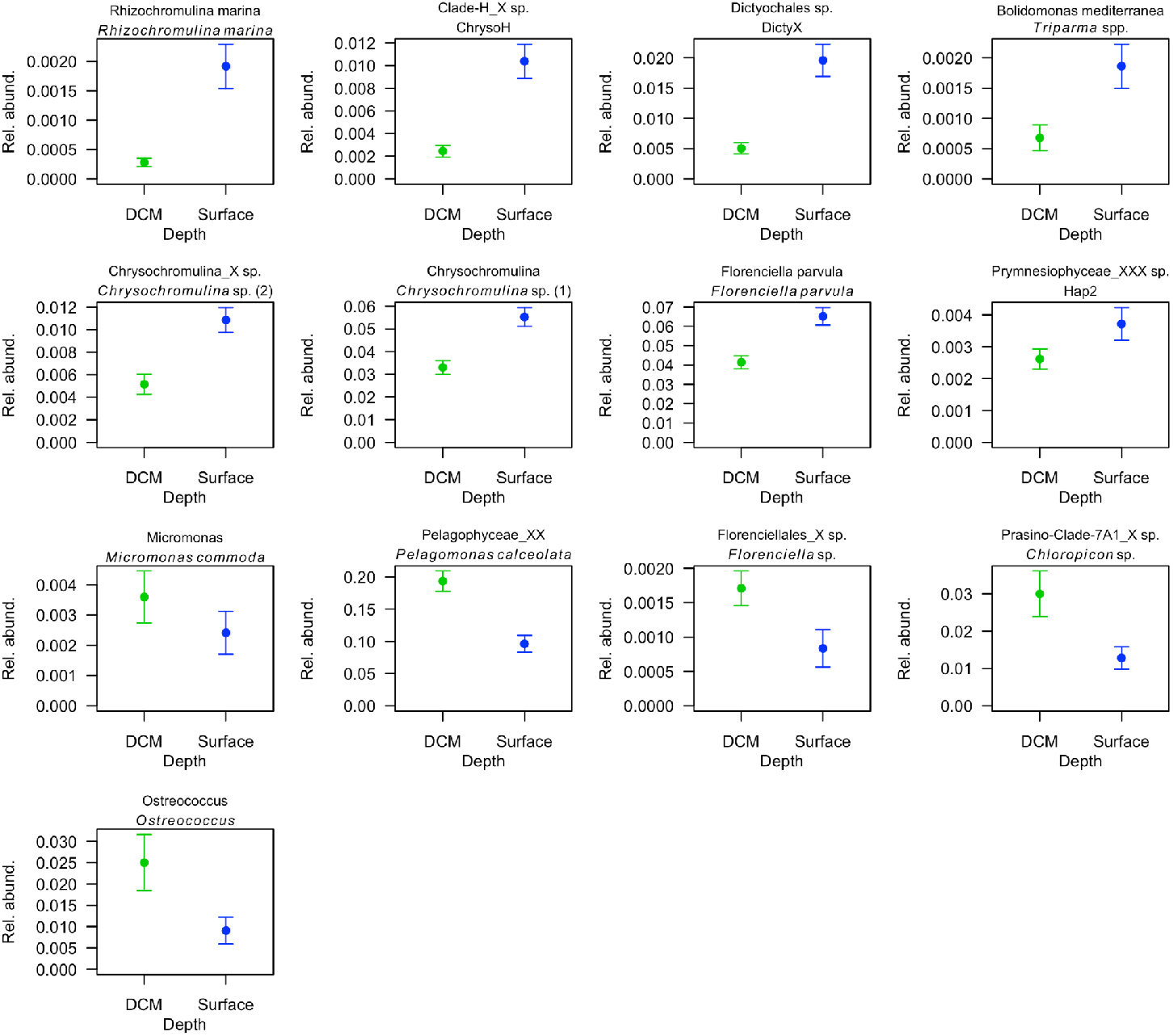
Relative abundance of 13 OTUs in deep chlorophyll maximum (DCM) and surface (3-7 m) samples. For each panel, the first title is the taxonomic annotation of the OTU in the *Tara* Oceans dataset, and the second title is the annotation of the corresponding isolate (see Materials and Methods; Table S2). Error bars are +/− one standard error of the mean.

**Figure S5.**
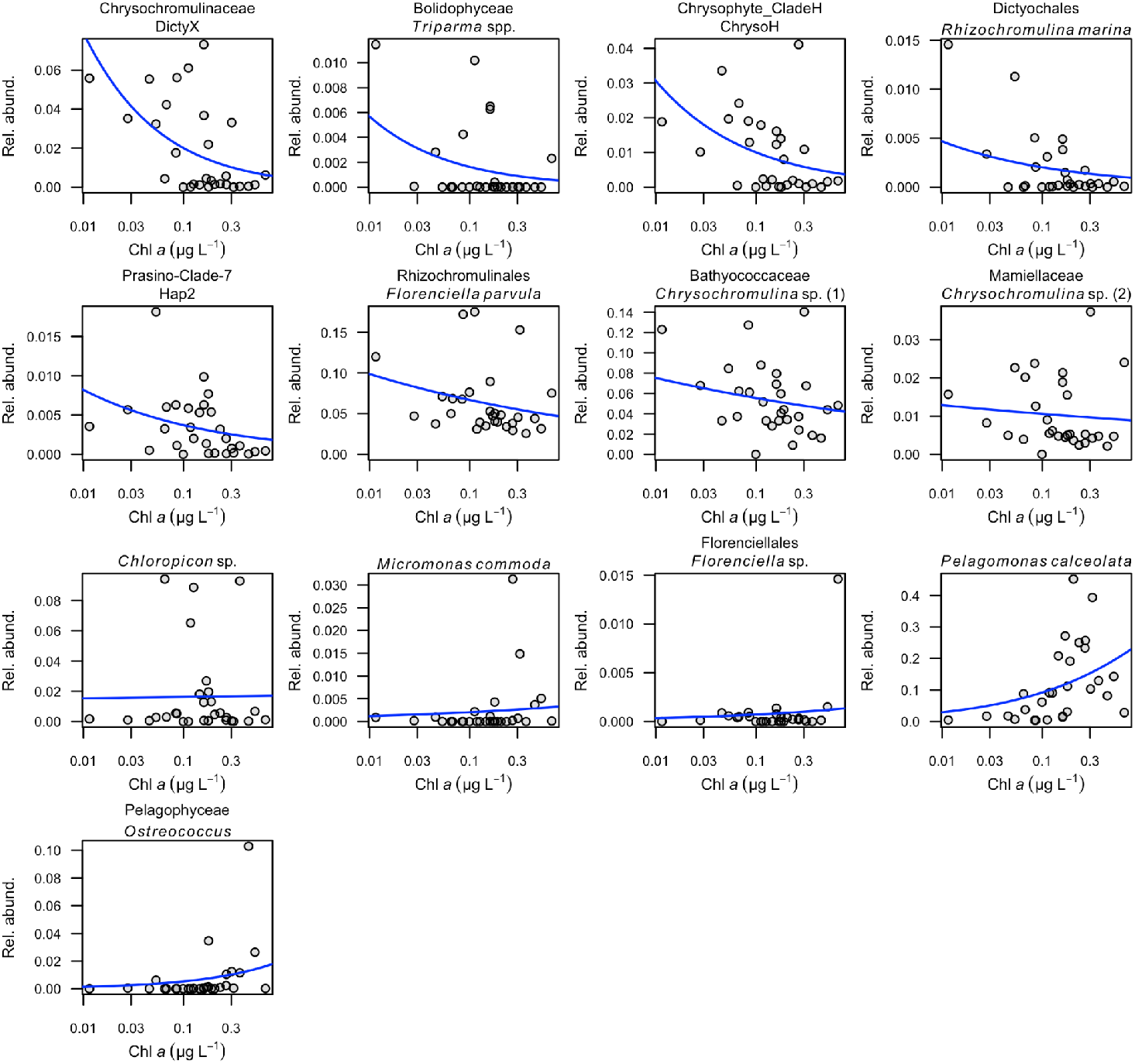
Relative abundance of 13 OTUs in surface (3-7 m) samples vs. chlorophyll *a* concentration determined by HPLC. For each panel, the first title is the taxonomic annotation of the OTU in the *Tara* Oceans dataset, and the second title is the annotation of the corresponding isolate (see Materials and Methods; Table S2). The curve is the fitted relationship from a beta-binomial GLMM with logit link function (see Materials and Methods).

**Figure S6.**
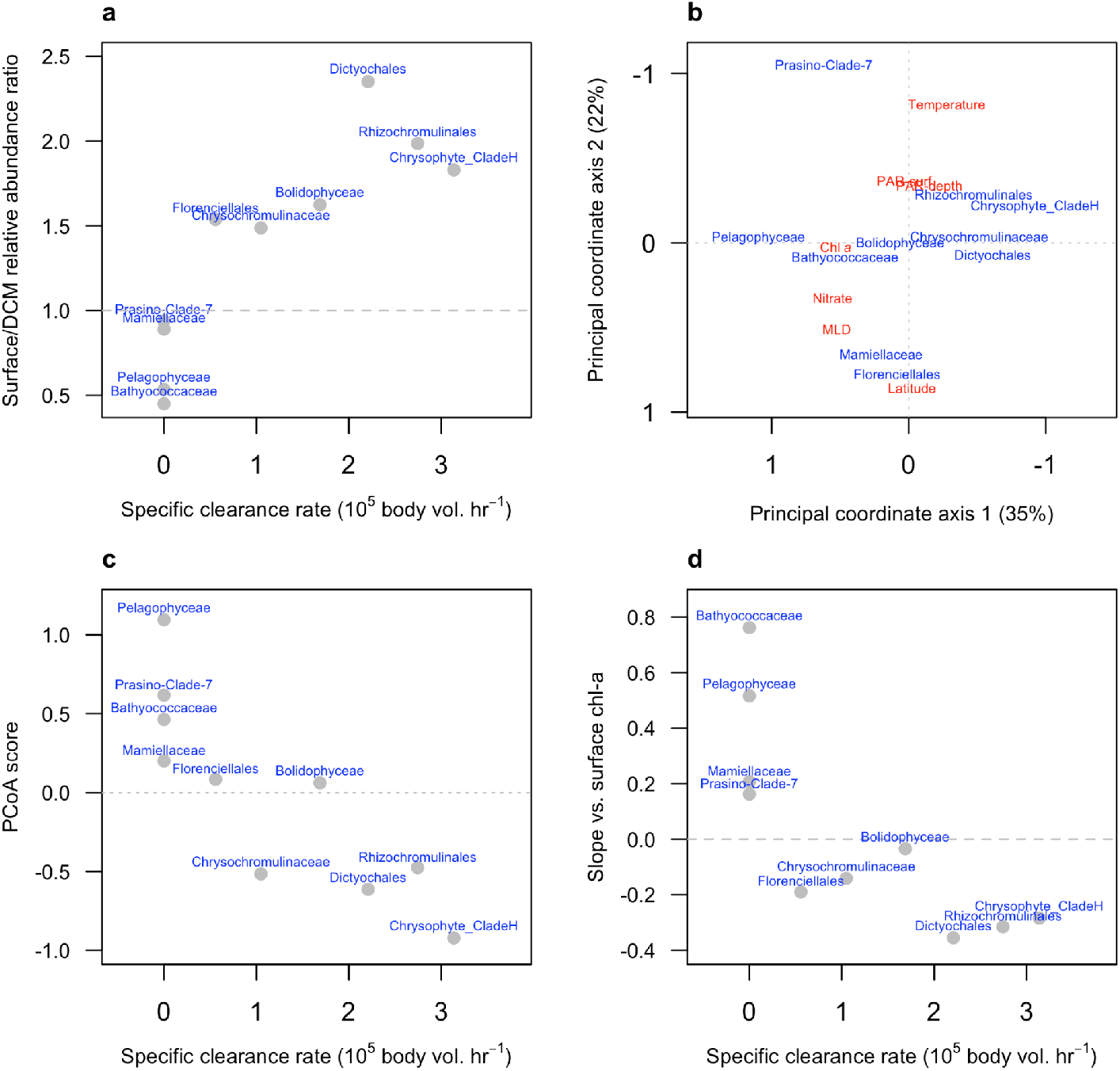
Relationship between environmental niches of broader clades and grazing ability of representative isolates. Class-, order-, or family-level clades, as annotated in the *Tara* Oceans metabarcode dataset, were chosen that include the OTUs analyzed in Fig. 2 (Table S2). Clades were chosen to utilize the broadest possible clades without lumping isolates from different genera (e.g., the three dictyochophyte genera are represented by the three different families to which they belong, while *Pelagomonas* is represented by the class Pelagophyceae). All reads within each taxonomic group were summed, and the relative abundance of the groups was analyzed using the same GLMM and PCoA approaches used for the OTUs (see Materials and Methods; Fig. 2). On average these groups account for 54% of all non-dinoflagellate phytoplankton reads in the 0.8-5 *μ*m size fraction. The response of the groups to environmental variables/axes was compared to the grazing ability (specific clearance rate) of the isolates belonging to those groups. For groups with multiple isolates (Florenciellales, Chrysochromulinaceae, Dictyochales, Bolidophyceae), grazing ability was averaged to yield a group mean.

**Figure S7.**
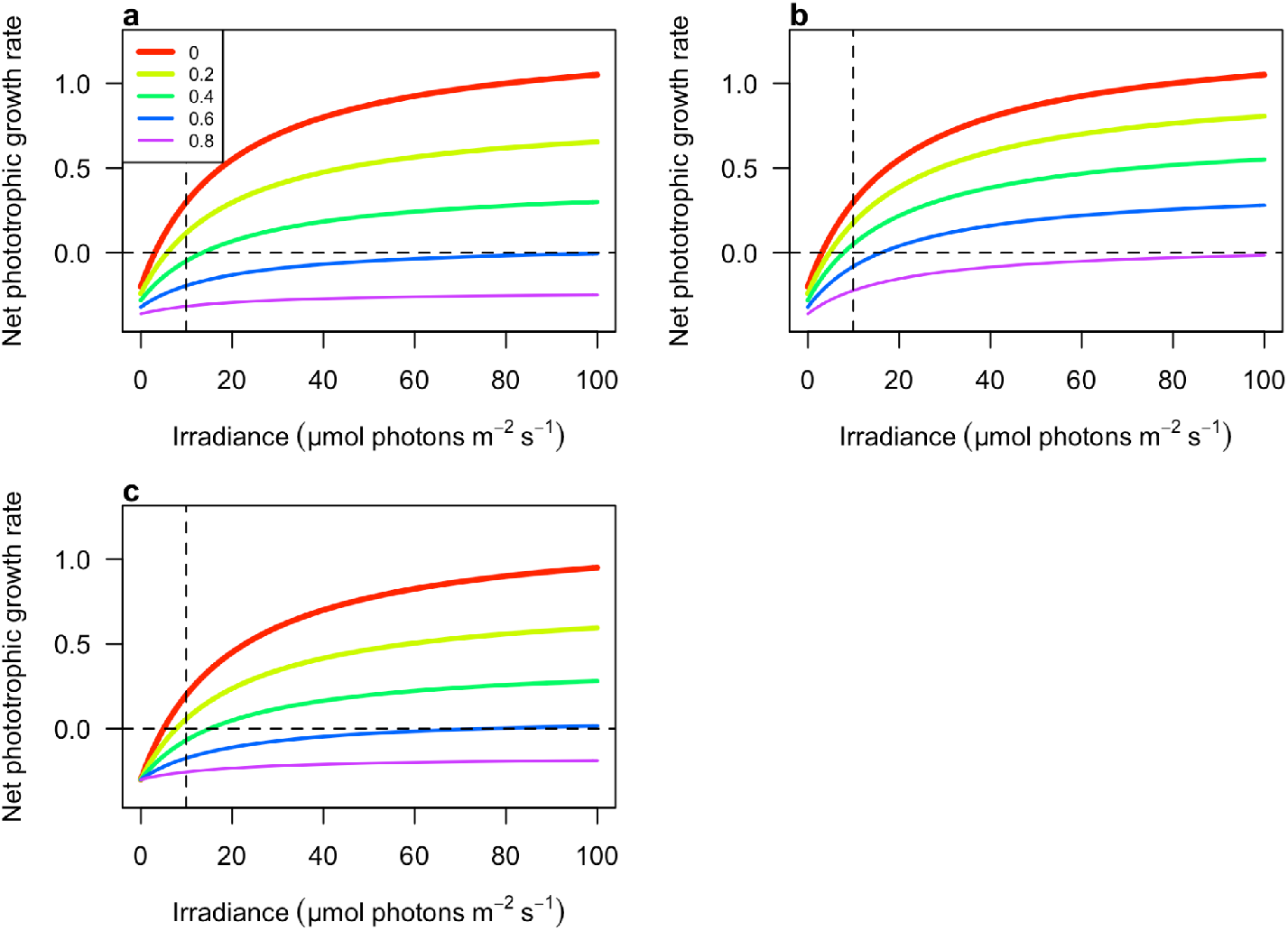
Illustration of the tradeoff scenarios used in the model of trophic strategy competition. All panels show net growth under phototrophic conditions vs. irradiance, for five example strategies ranging from autotrophic (*x* = 0) to mostly heterotrophic (*x* = 0.8). (a) Growth curves for the tradeoff parameters used in Fig. 3 (*ϕ* = 1.5, *m*_0_ = 0.2, *m*_1_ = 0.2). In this scenario the autotroph can grow readily at 10 *μ*mol photons m^-2^ s^-1^ (vertical dashed line), while most mixotrophs cannot, and relatively heterotrophic mixotrophs also cannot grow under high light (100 *μ*mol photons m^-2^ s^-1^). These patterns approximate those found in the experiments (Fig. 1). (b) Growth curves under a weaker tradeoff assumption (Fig. S9; *ϕ* = 0.8, *m*_0_ = 0.2, *m*_1_ = 0.2). In this scenario mixotrophs can grow more readily under low light, as well as high light. (c) Growth curves with no assumed correlation between phagotrophy and maintenance respiration (Fig. S10; *ϕ* = 1.5, *m*_0_ = 0.3, *m*_1_ = 0). In this scenario the autotrophs differ most from panel (a), growing more slowly under low light.

**Figure S8.**
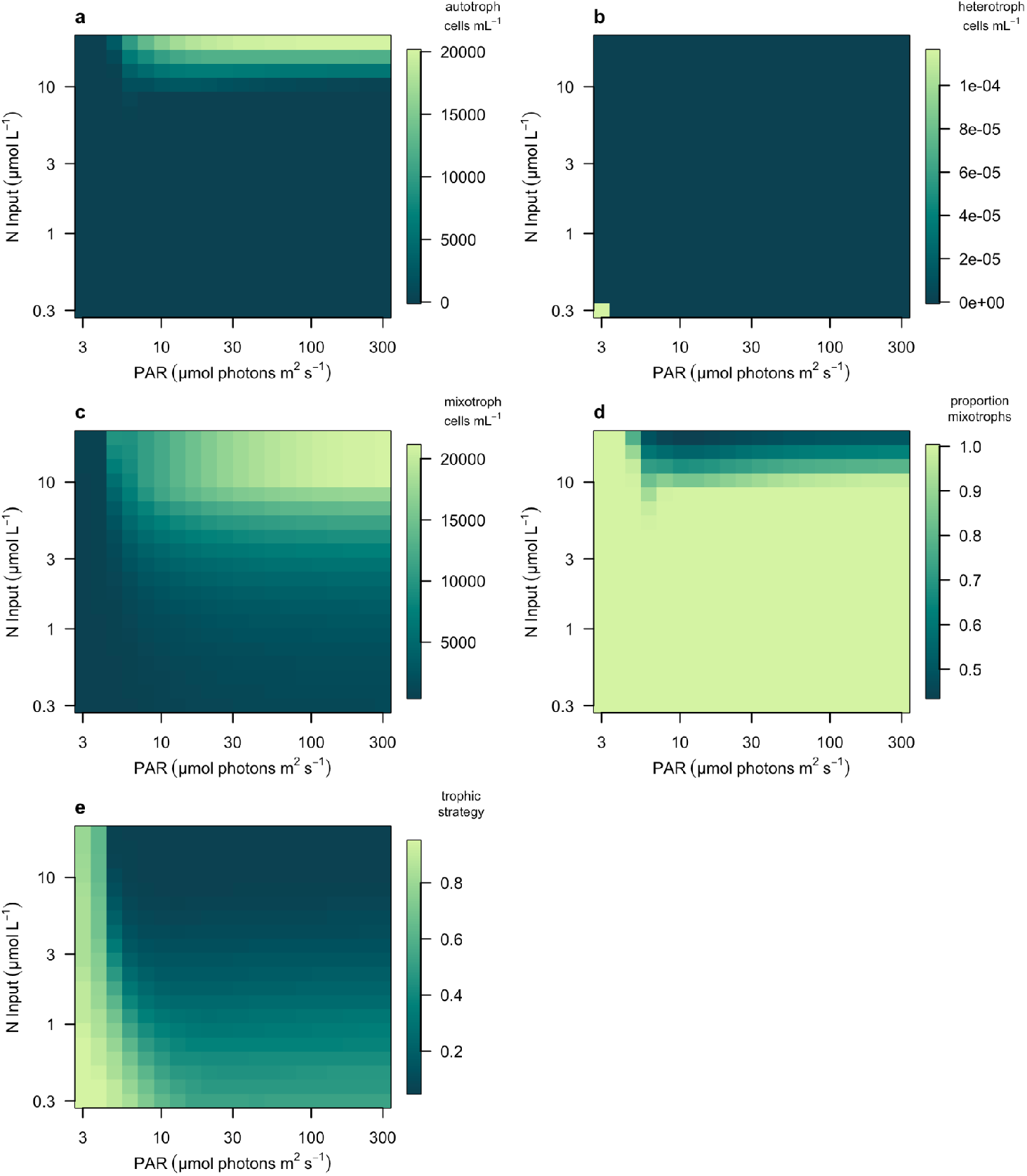
Modeled trophic strategies vs. gradients of nitrogen input and irradiance, assuming a weaker tradeoff between phototrophic and phagotrophic functions (*ϕ* = 0.8, *m*_0_ = 0.2, *m*_1_ = 0.2). (a) Concentration of autotrophs. (b) Concentration of heterotrophs. (c) Concentration of mixotrophs. (d) Mixotrophs as a proportion of all phytoplankton. (e) Trophic strategy of the persisting mixotroph population, where strategy is the parameter *x* (see Materials and Methods) which ranges from 0 (autotroph) to 1 (heterotroph).

**Figure S9.**
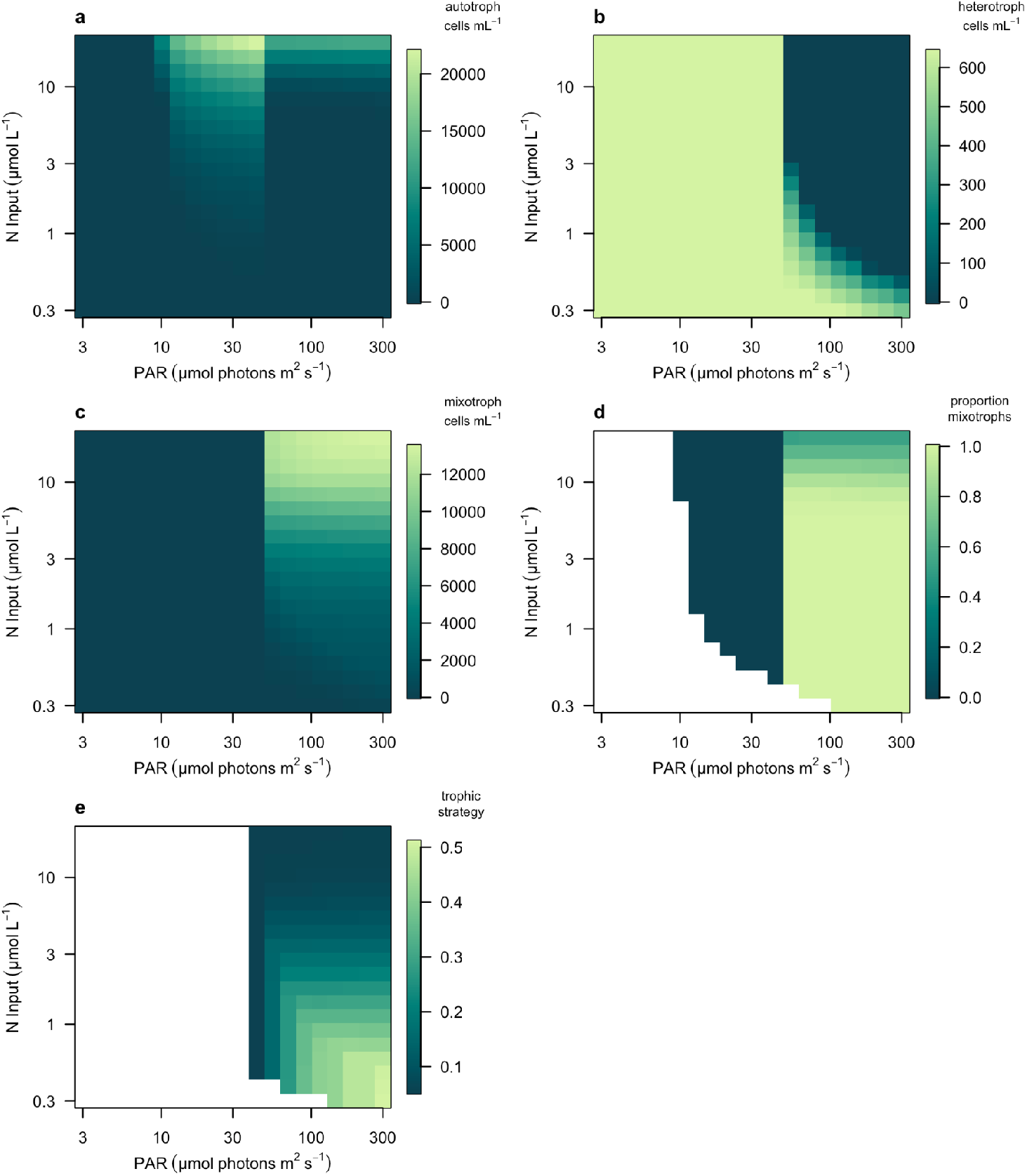
Modeled trophic strategies vs. gradients of nitrogen input and irradiance, assuming no correlation between phagotrophy and maintenance respiration (*ϕ* = 1.5, *m*_0_ = 0.3, *m*_1_ = 0). (a) Concentration of autotrophs. (b) Concentration of heterotrophs. (c) Concentration of mixotrophs. (d) Mixotrophs as a proportion of all phytoplankton. (e) Trophic strategy of the persisting mixotroph population, where strategy is the parameter *x* (see Materials and Methods) which ranges from 0 (autotroph) to 1 (heterotroph). Regions in white denote where phytoplankton did not persist (panel a) or mixotrophs did not persist (panel b).

## Supplementary Tables

**Table S1.** Strains used in the phototrophic and phagotrophic growth experiments. For strains from undescribed genera abbreviations used in the main text are listed (ChloraX, ChrysoH - see Materials and Methods). All strains were isolated from the euphotic zone at Station ALOHA (22° 45’ N, 158° 00’ W) between 2011-2019. ‘HL’ and ‘LL’ growth rate are high light and low light, respectively, as described in the main text. Numbers are means from 1-4 replicate growth curves (Fig. S3). ‘Phagotrophic growth’ describes whether the strain can grow when fed *Prochlorococcus* as the only added nitrogen source, under high light conditions (Fig. S2, (*17*)).

| Class | Species | Strain ID | Accession # | HL growth rate | LL growth rate | Phagotrophic growth |
| --- | --- | --- | --- | --- | --- | --- |
| Chlorarachniophyceae | Chlorax | UHM2000 | MZ611732 | 0.78 | 0 | y |
| Chloropicophyceae | *Chloropicon* sp. | UHM1200 | OM521953 | 1.1 | 0.25 | n |
| Prymnesiophyceae | *Chrysochromulina* sp. (1) | UHM4110 | MZ611704 | 0.52 | 0 | y |
| Prymnesiophyceae | *Chrysochromulina* sp. (2) | UHM4100 | MZ611712 | 0.67 | 0 | y |
| Chrysophyceae | ChrysoH | UHM3501 | MZ611720 | 0 | 0 | y |
| Dictyochophyceae | *Florenciella* sp. (1) | UHM3020 | MN615710 | 2.4 | 0 | y |
| Dictyochophyceae | *Florenciella* sp. (2) | UHM3021 | MN615711 | 1.4 | 0 | y |
| Mamiellophyceae | *Micromonas commoda* | UHM1080 | OM521954 | 1.5 | 0.27 | n |
| Mamiellophyceae | *Ostreococcus* sp. | UHM1000 | OM521955 | 0.88 | 0.24 | n |
| Pelagophyceae | *Pelagomonas calceolata* | UHM3400 | OM521952 | 1.4 | 0.34 | n |
| Bolidophyceae | *Triparma mediterranea* | UHM3600 | MZ611730 | 0.46 | 0.28 | y |

**Table S2.** Strains used for the comparison of grazing ability and environmental niches. Class and species assignments are putative based on 18S phylogenies. For strains from undescribed genera the abbreviations used in the main text are listed (ChrysoH, DictyX, Hap2 - see Materials and Methods). Strain ID = reference number in our culture collection. All strains were isolated from the euphotic zone at Station ALOHA (22° 45’ N, 158° 00’ W) between 2011-2019. Strains were matched to *Tara* Oceans metabarcode OTUs (see Materials and Methods); ‘*Tara* species’ is the lowest level OTU taxonomic annotation in the *Tara* dataset; *‘Tara* ID’ lists the identifier of the 18S rDNA V9 reference sequence of the OTU (http://taraoceans.sb-roscoff.fr/EukDiv/). ‘Specific clearance rate’ is the mean body volume-specific clearance rate (10^5^ body volumes hr^-1^), for all strains matched to the corresponding *Tara* Oceans OTU. Clearance rate data is taken from Li et al. (*17*). Clearance rate of the strains that did not show phagotrophic growth (Table S1) is denoted ‘NA’.

| Class | Species | Strain ID | *Tara* species | *Tara* ID | Specific clearance rate |
| --- | --- | --- | --- | --- | --- |
| Bolidophyceae | *Triparma* spp. | UHM3600,<br>UHM3610 | Bolidomonas mediterranea | 239078 | 1.7 |
| Chrysophyceae | ChrysoH | UHM3500,<br>UHM3501,<br>UHM3502 | Clade-H_X sp. | 115485 | 3.1 |
| Dictyochophyceae | *Florenciella* sp. | UHM3023,<br>UHM3029,<br>UHM3020,<br>UHM3027,<br>UHM3021 | Florenciellales_X sp. | 73377 | 0.27 |
| Dictyochophyceae | *Florenciella parvula* | UHM3011,<br>UHM3012,<br>UHM3010 | *Florenciella parvula* | 146146 | 0.85 |
| Dictyochophyceae | *Rhizochromulina marina* | UHM3070,<br>UHM3073,<br>UHM3072,<br>UHM3071 | *Rhizochromulina marina* | 67389 | 2.7 |
| Dictyochophyceae | DictyX | UHM3050,<br>UHM3060,<br>UHM3052,<br>UHM3051,<br>UHM3054 | Dictyochales sp. | 46831 | 2.2 |
| Prymnesiophyceae | *Chrysochromulina* sp. (2) | UHM4100 | Chrysochromulina | 11632 | 0.52 |
| Prymnesiophyceae | *Chrysochromulina* sp. (1) | UHM4110 | Chrysochromulina_X sp. | 178099 | 1.6 |
| Prymnesiophyceae | Hap2 | UHM4150 | Prymnesiophyceae_XXX sp. | 162435 | 1.2 |
| Mamiellophyceae | *Ostreococcus* sp. | UHM1000 | Ostreococcus | 153479 | NA |
| Mamiellophyceae | *Micromonas commoda* | UHM1080 | Micromonas | 86716 | NA |
| Chloropicophyceae | *Chloropicon* sp. | UHM1200 | Prasino-Clade-7A1_X sp. | 40975,<br>151962 | NA |
| Pelagophyceae | *Pelagomonas calceolata* | UHM3400 | Pelagophyceae_XX | 161224 | NA |

**Table S3.**
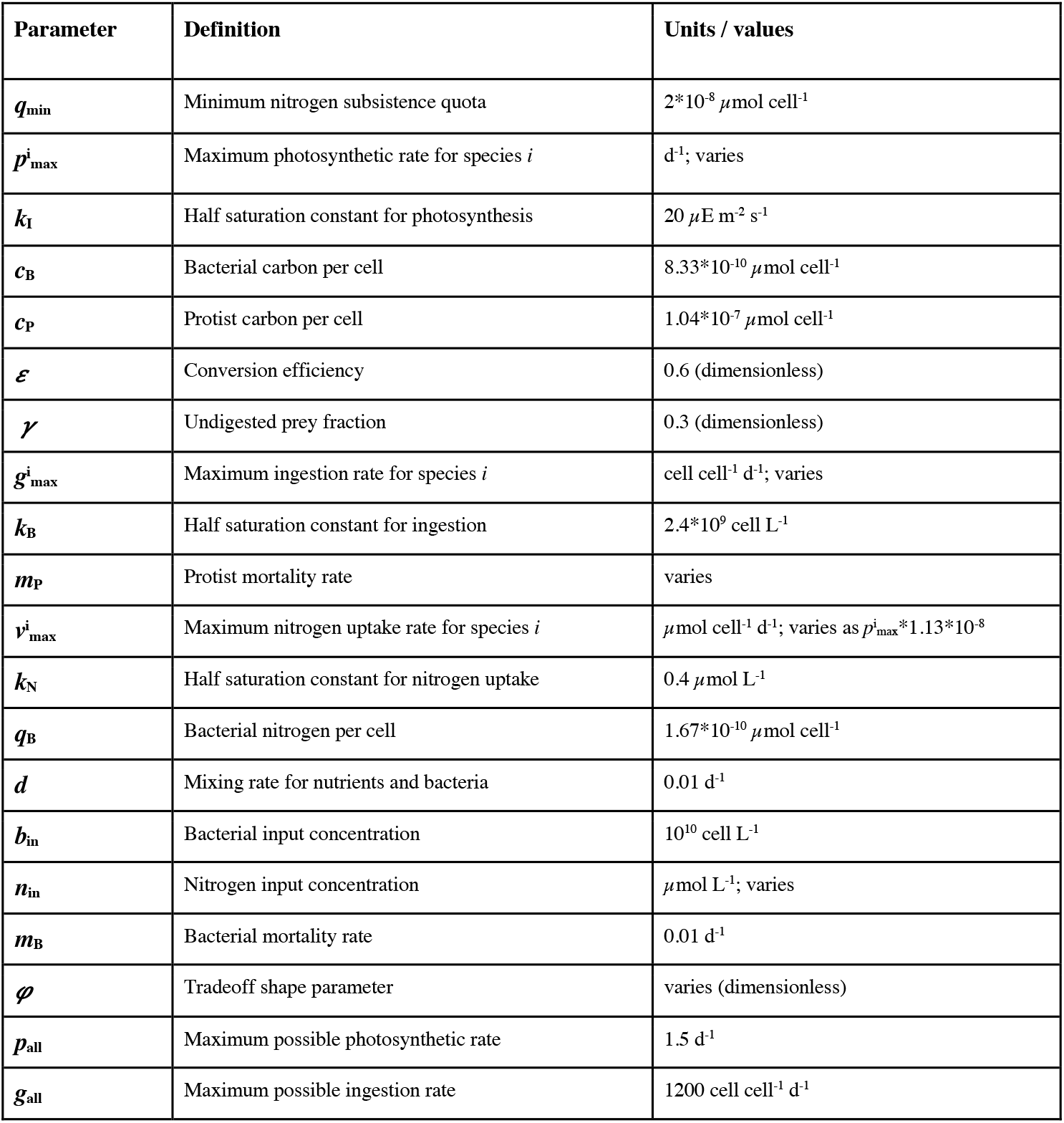
Model parameter definitions and values. Parameters are assigned values typical for small unicellular protists, based on known allometric relationships (*27, 53, 54*).

